# HCMV glycoprotein B nucleoside-modified mRNA vaccine elicits antibody responses with greater durability and breadth than MF59-adjuvanted gB protein immunization

**DOI:** 10.1101/797894

**Authors:** Cody S. Nelson, Jennifer A. Jenks, Norbert Pardi, Matthew Goodwin, Hunter Roark, Whitney Edwards, Jason S. McLellan, Justin Pollara, Drew Weissman, Sallie R. Permar

## Abstract

A vaccine to prevent maternal acquisition of human cytomegalovirus (HCMV) during pregnancy is a primary strategy to reduce the incidence of congenital disease. Similarly, vaccination of transplant recipients against HCMV has been proposed to prevent transplant-associated HCMV morbidity. The MF59-adjuvanted glycoprotein B protein subunit vaccine (gB/MF59) is the most efficacious tested to-date for both indications. We previously identified that gB/MF59 vaccination elicited poor neutralizing antibody responses and an immunodominant response against gB antigenic domain 3 (AD-3). Thus, we sought to test novel gB vaccines to improve functional antibody responses and reduce AD-3 immunodominance. Groups of juvenile New Zealand White rabbits were administered 3 sequential doses of full-length gB protein with an MF59-like squalene adjuvant (analogous to clinically-tested vaccine), gB ectodomain protein (lacking AD-3) with squalene adjuvant, or lipid nanoparticle (LNP)-packaged nucleoside-modified mRNA encoding full-length gB. The AD-3 immunodominant IgG response following human gB/MF59 vaccination was closely mimicked in rabbits, with 78% of binding antibodies directed against this region in the full-length gB protein group compared to 1% and 46% in the ectodomain and mRNA-LNP-vaccinated groups, respectively. All vaccines were highly immunogenic with similar kinetics and comparable peak gB-binding and functional antibody responses. Although gB ectodomain subunit vaccination reduced targeting of non-neutralizing epitope AD-3, it did not improve vaccine-elicited neutralizing or non-neutralizing antibody functions. gB nucleoside-modified mRNA-LNP-immunized rabbits exhibited enhanced durability of IgG binding to soluble and cell membrane-associated gB protein as well as HCMV-neutralizing function. Furthermore, the gB mRNA-LNP vaccine enhanced breadth of IgG binding responses against discrete gB peptide residues. Finally, low-magnitude gB-specific T cell activity was observed in the full-length gB protein and mRNA-LNP vaccine groups, though not in ectodomain-vaccinated rabbits. Altogether, these data suggest that the gB mRNA-LNP vaccine candidate, aiming to improve upon the partial efficacy of gB/MF59 vaccination, should be further evaluated in preclinical models.

**Author summary:** Human cytomegalovirus (HCMV) is the most common infectious cause of infant birth defects, resulting in permanent neurologic disability for one newborn child every hour in the United States. Furthermore, this virus causes significant morbidity and mortality in immune-suppressed transplant recipients. After more than a half century of research and development, we remain without a clinically-licensed vaccine or therapeutic to reduce the burden of HCMV-associated disease. In this study, we sought to improve upon the glycoprotein B protein vaccine (gB/MF59), the most efficacious HCMV vaccine evaluated in clinical trial, via targeted modifications to either the protein structure or vaccine formulation. An attempt to alter the protein structure to focus the immune response on vulnerable epitopes (‘gB ectodomain’) had little effect on the quality or function of the vaccine-elicited antibodies. However, a novel vaccine platform, nucleoside-modified mRNA formulated in lipid nanoparticles, increased the durability and breadth of vaccine-elicited immune responses. We propose that an mRNA-based gB vaccine may ultimately prove more efficacious than the gB/MF59 vaccine and should be further evaluated for its ability to elicit antiviral immune factors that can prevent both infant and transplant-associated disease caused by HCMV infection.

## Introduction

Human cytomegalovirus (HCMV) impacts 1 in 150 live born infants, making this pathogen the most common cause of congenital infection worldwide [1, 2]. Approximately 20% of infants infected with HCMV *in utero* will develop long-term sequelae including microcephaly, intrauterine growth restriction, hearing/vision loss, or neurodevelopmental delay [3, 4]. Furthermore, HCMV is the most prevalent infection among solid organ and hematopoietic stem cell transplant recipients, causing end-organ disease such as gastroenteritis, pneumonitis, or hepatitis and potentially predisposing these individuals to allograft rejection and/or failure [5, 6]. However, we remain without a vaccine or immunotherapeutic intervention to reduce the burden of disease among newborn children and transplant recipients.

A variety of vaccine platforms and formulations have been trialed for the prevention of both congenital (reviewed in [7]) and transplant-associated HCMV disease (reviewed in [8]), of which the most efficacious has been the glycoprotein B (gB) subunit vaccine administered with MF59 squalene adjuvant [9]. gB is the viral fusogen and is essential for entry into all cell types [10], including placental trophoblast progenitor cells [11]. Furthermore, gB is highly-expressed and an immune-dominant target following natural infection, making this protein an attractive target for vaccination. gB/MF59 subunit vaccination demonstrated moderate (∼50%) efficacy in blocking HCMV infection and host seroconversion in populations of HCMV-seronegative postpartum [12] and adolescent women [13]. Furthermore, in transplant recipients this vaccine protected against HCMV viremia and reduced the clinical need for antiviral treatment [14].

Analysis of samples obtained from both postpartum and transplant-recipient gB vaccinees revealed two key observations regarding the target and function of gB-elicited antibody responses that inform our understanding of the partial vaccine efficacy. First, we identified that vaccination elicited an extraordinarily robust response against antigenic domain 3 (AD-3), a cytosolic non-neutralizing epitope in the C-terminal region of the protein [15]. Second, we noted that gB-specific antibodies elicited in postpartum women and transplant-recipients were predominantly non-neutralizing, suggesting that the mechanism of partial protection against viral acquisition was not the induction of neutralizing antibodies [15, 16]. However, the protective non-neutralizing function remains unclear: we identified that gB/MF59 vaccinees had high-magnitude viral phagocytosis activity, though magnitude was not associated with infection status [15]. These results led us to hypothesize that we might improve upon the gB/MF59 vaccine through rational design of novel immunogens that minimize responses against the intracellular to gB AD-3 epitope.

Here we present an investigation into the immunogenicity of two novel gB vaccines aiming to reduce exposure to AD-3: a truncated gB protein subunit vaccine (lacking the AD-3 epitope) administered with MF59-like squalene adjuvant AddaVax (gB ectodomain vaccine) and a gB nucleoside-modified mRNA vaccine packaged in lipid nanoparticles (gB mRNA-LNP vaccine). New Zealand White rabbits were selected for this study because we previously identified that neutralizing and non-neutralizing antibodies are elicited in rabbits by vaccination, and that F_c_ receptor-independent and dependent effector functions can be measured *in vitro* [17]. Three groups of rabbits were vaccinated with either: 1) full-length gB + AddaVax (immunogen from gB/MF59 vaccine trial; ‘gB FL’), 2) gB ectodomain + AddaVax (‘gB ecto’), or 3) gB mRNA-LNP (‘gB mRNA). We anticipated that gB ectodomain and gB mRNA-LNP vaccines would have reduced targeting of AD-3 and enhanced neutralizing and/or non-neutralizing function, resulting in a superior vaccine that might be deployed to prevent both congenital and transplant-associated HCMV disease.

## Results

### IgG binding to soluble and cell-associated gB

We first assessed the ability of IgG elicited by all three vaccines (**Figure 1;** gB FL, gB ecto, and gB mRNA) to bind soluble full-length gB (**Figure 2A**) and gB ectodomain proteins (**Figure 2B**) by BAMA, as well as cell-associated gB on the surface of gB-transfected cells by flow cytometry (**Figure 2C**). All vaccines were highly immunogenic, with similar peak immunogenicity (10 weeks) binding magnitude to both soluble and cell-associated gB. However, gB mRNA-immunized rabbits had enhanced binding to both soluble and cell-associated gB at the time of animal necropsy (20 weeks), indicating superior durability of the mRNA vaccine-elicited antibody responses compared to gB FL. This distinction was most pronounced for binding to cell-associated gB (median % PE+ cells at 20 weeks: mRNA = 29.1% vs FL = 18.8%, p=0.01, Kruskal-Wallis + *post hoc* Mann-Whitney U test). Additionally, we evaluated the avidity of vaccine-elicited IgG responses by plate-based, urea wash ELISA (**Figure 2D**). We noted slightly reduced median RAI (relative avidity index, measured against soluble gB FL protein) in gB mRNA-immunized rabbits, though the difference was not statistically significant.

**Figure 1.**
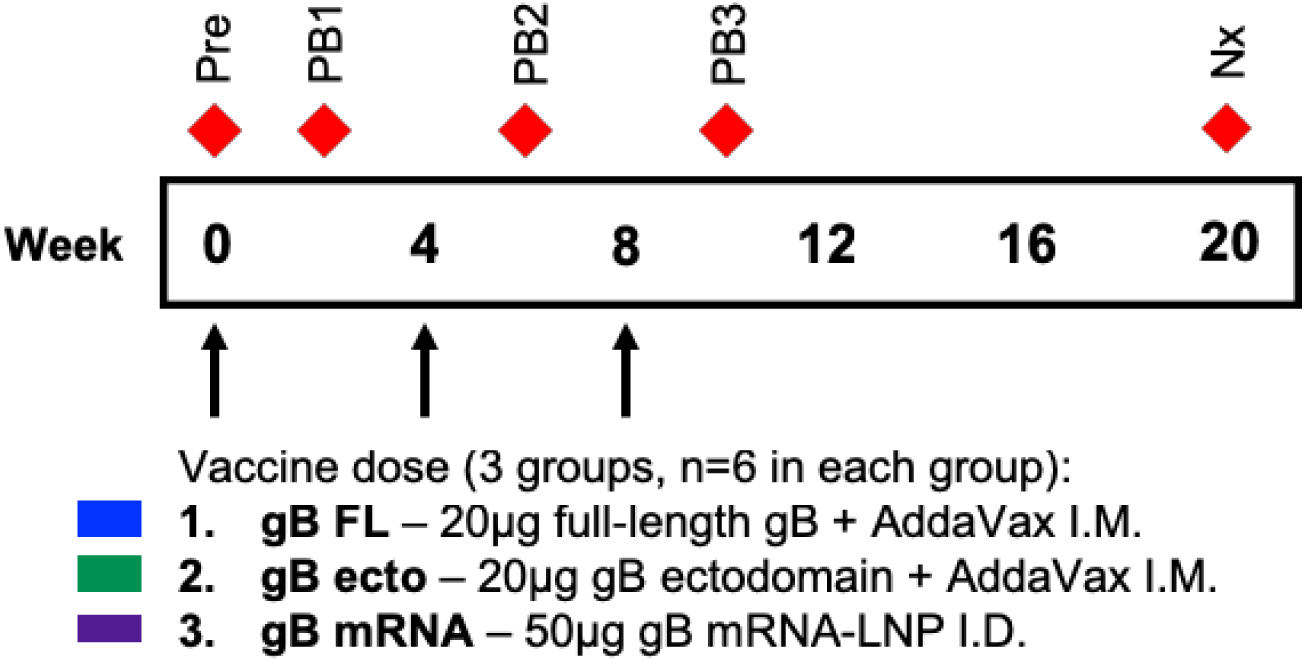
Vaccination and sampling timeline. 6At 0, 4, and 8 weeks, juvenile New Zealand White rabbits were administered 50μg doses of either full-length gB protein + AddaVax intramuscularly (blue), gB ectodomain protein + AddaVax intramuscularly (green), or lipid nanoparticle-packaged gB mRNA intradermally (purple). Blood was sampled at the following timepoints indicated by red diamonds: preimmune (0 weeks; ‘Pre’), post boost 1 (2 weeks, ‘PB1’), post boost 2 (6 weeks, ‘PB2’), post boost 3 (10 weeks, ‘PB3’), and necropsy (20 weeks, ‘Nx’).

**Figure 2.**
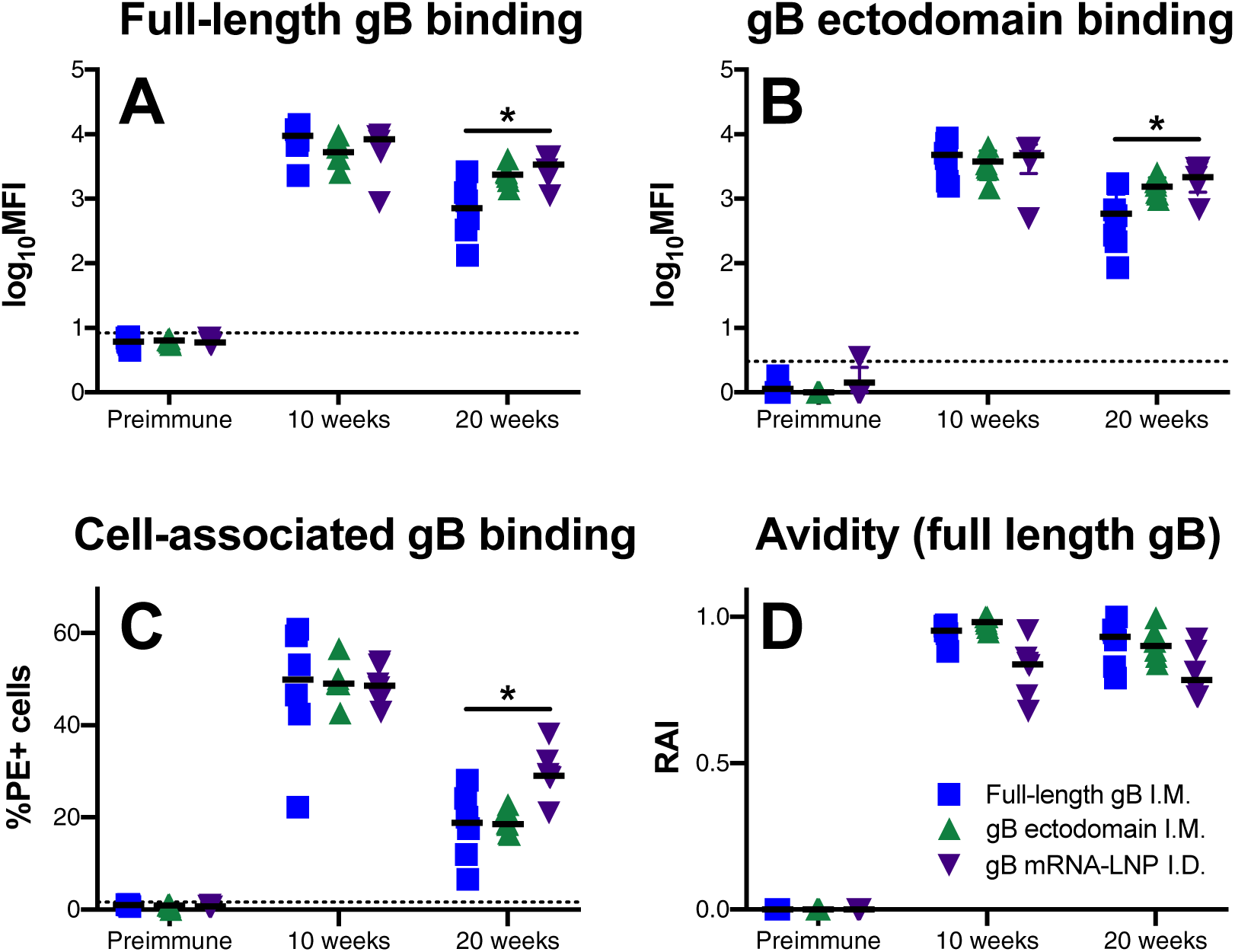
gB mRNA-LNP vaccination elicits more durable binding antibody responses against soluble and cell-associated gB. IgG binding to full-length gB (A) and gB ectodomain(B) proteins were assessed by BAMA. IgG binding to gB-transfected cells (C) was measured by flow cytometry. gB binding avidity was assessed against full-length gB using urea-wash ELISA. All proteins were strain-matched (Towne). IgG responses for full-length gB vaccinees are shown in blue, gB ectodomain in green, and gB mRNA-LNP in purple. Binding responses were assessed for preimmune, 10 week (post boost 3, peak immunogenicity), and 20 week (necropsy) timepoints. Data points represent individual animals, with the line designating the median. Dotted black line indicates the mean preimmune response + 2 standard deviations. *p<0.05, Kruskal-Wallis + *post hoc* Mann-Whitney U test.

### Linear gB epitope binding

To identify the epitope specificity and breadth of IgG responses elicited by each vaccine, we utilized a peptide microarray library consisting of 15-mers overlapping each subsequent peptide by 10 residues and spanning the entire gB ORF (Towne strain) (**Figure 3A**). We observed that rabbits administered the gB FL vaccine had a nearly identical AD-3 epitope immunodominance to that observed in human gB/MF59 vaccinees [18], with 77% of the peptide-binding IgG response directed against this singular region (vs. 78% in human vaccinees). AD-3 linear peptide binding was dramatically reduced in gB ecto (<1%) and gB mRNA (46%) groups. However, AD-3 remained the dominant response in gB FL and gB mRNA groups, while the furin cleavage site was dominant for gB ecto (no AD-3 in the immunogen) (**Figure S1**). Furthermore, gB mRNA vaccinated rabbits had slightly reduced total peptide binding compared to gB FL (**Figure 3B**; median peptide-binding MFI sum: FL = 96,629, mRNA = 48,051, p=ns, Kruskal-Wallis + *post hoc* Mann-Whitney U test), though both gB FL and gB mRNA had greater total peptide binding than the gB ecto group (both p<0.05, Kruskal-Wallis + *post hoc* Mann-Whitney U test). Importantly, there was enhanced breadth of peptide-binding responses in gB mRNA vaccinated rabbits compared to both gB FL and gB ecto groups (**Figure 3D**; median number of discrete peptides bound: FL = 44.5, ecto = 28.5, mRNA = 85, both p<0.05, Kruskal-Wallis + *post hoc* Mann-Whitney U test). Of note, the pattern of linear peptide binding and breadth of mRNA-immunized rabbits appears analogous to that elicited by natural HCMV infection [18].

**Figure 3.**
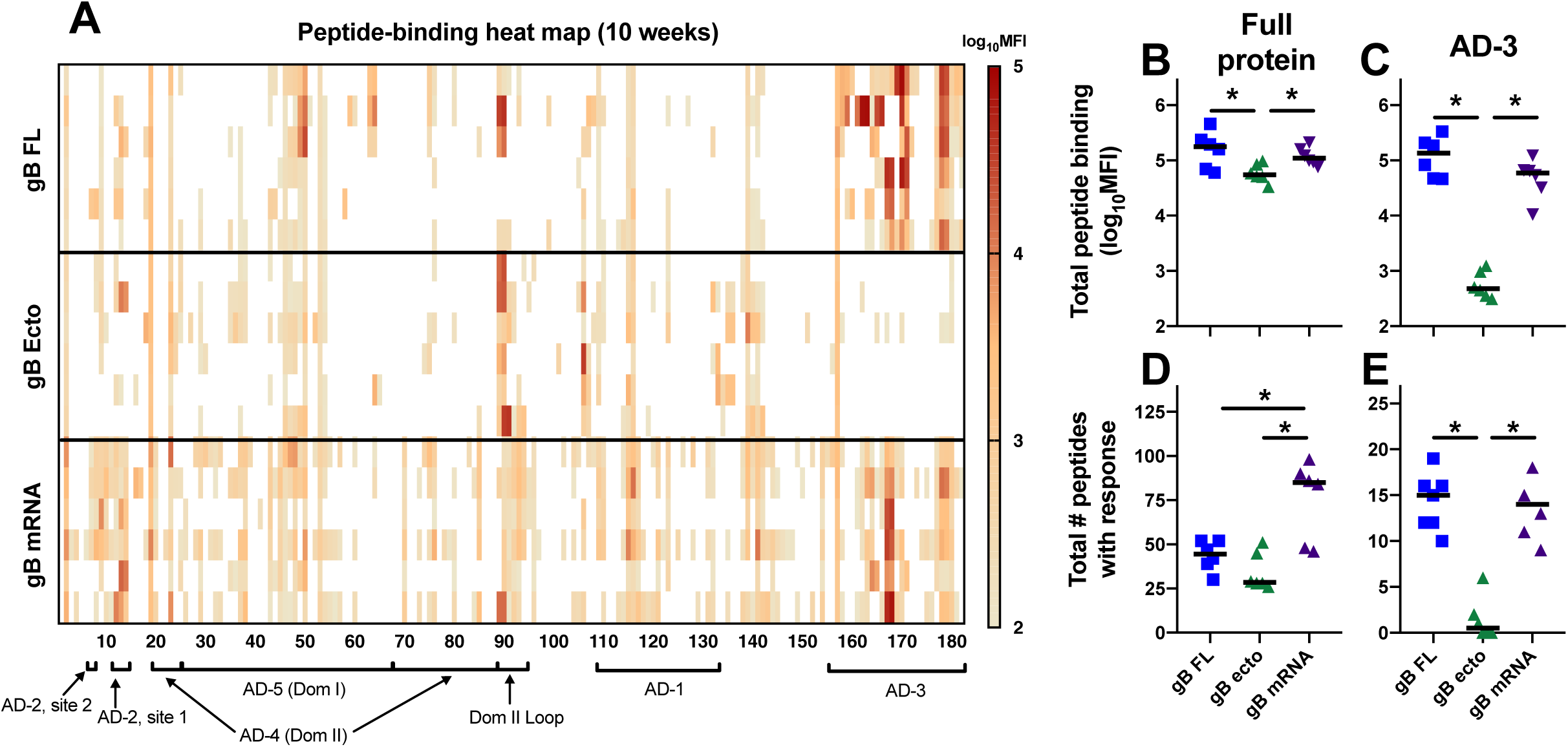
gB mRNA-LNP vaccination reduced AD-3 immunodominance, yet enhances breadth of linear peptide binding IgG response. (A) The binding magnitude of rabbit antibodies at week 10 (post boost 3, peak immunogenicity) were assessed against a 15-mer peptide library spanning the entire Towne gB ORF (180 unique peptides). Each row indicates a single rabbit. Peptides corresponding to distinct gB antigenic domains are indicated along the x-axis. (B,C) The sum of total peptide-binding MFI to both the full gB protein (B) and those peptide corresponding to the AD-3 epitope (C). (D,E) Number of unique peptides with a binding response >100 MFI for the full gB protein (C) and AD-3 epitope (E). Responses for full-length gB vaccinees are shown in blue, gB ectodomain in green, and gB mRNA-LNP in purple. Data points represent individual animals, with the line designating the median. *p<0.05, Kruskal-Wallis + *post hoc* Mann-Whitney U test.

### Vaccine-elicited IgG binding to neutralizing epitopes and HCMV neutralization

We next examined IgG binding to regions of gB known to be targeted by neutralizing antibodies Domain 1 (AD-4), Domain 1+2 (AD-4 + AD-5), AD-1, and AD-2 (**Figure 4A-D**). Notably, we did not identify vaccine-elicited antibodies against AD-2 in any of the 3 vaccine groups (**Figure 4D**), an epitope known to be the target of potently-neutralizing gB-specific antibodies associated with reduced viremia in transplant recipients [19] and protection against congenital transmission [20]. Furthermore, low responses were noted against AD-1 (**Figure 4C**). The kinetics of vaccine-elicited IgG binding against Domain 1 as well as Domain 1+2 mirrored those of binding to soluble gB protein (**Figure 2**), However, at 20 weeks both gB ecto and gB mRNA vaccinated rabbits had enhanced binding against Domain 1 compared to gB FL (median Domain 1 MFI at 20 weeks: FL = 571, ecto = 2,343, mRNA = 2,730, p<0.05 for both, Kruskal-Wallis + *post hoc* Mann-Whitney U test) (**Figure 4A**).

**Figure 4.**
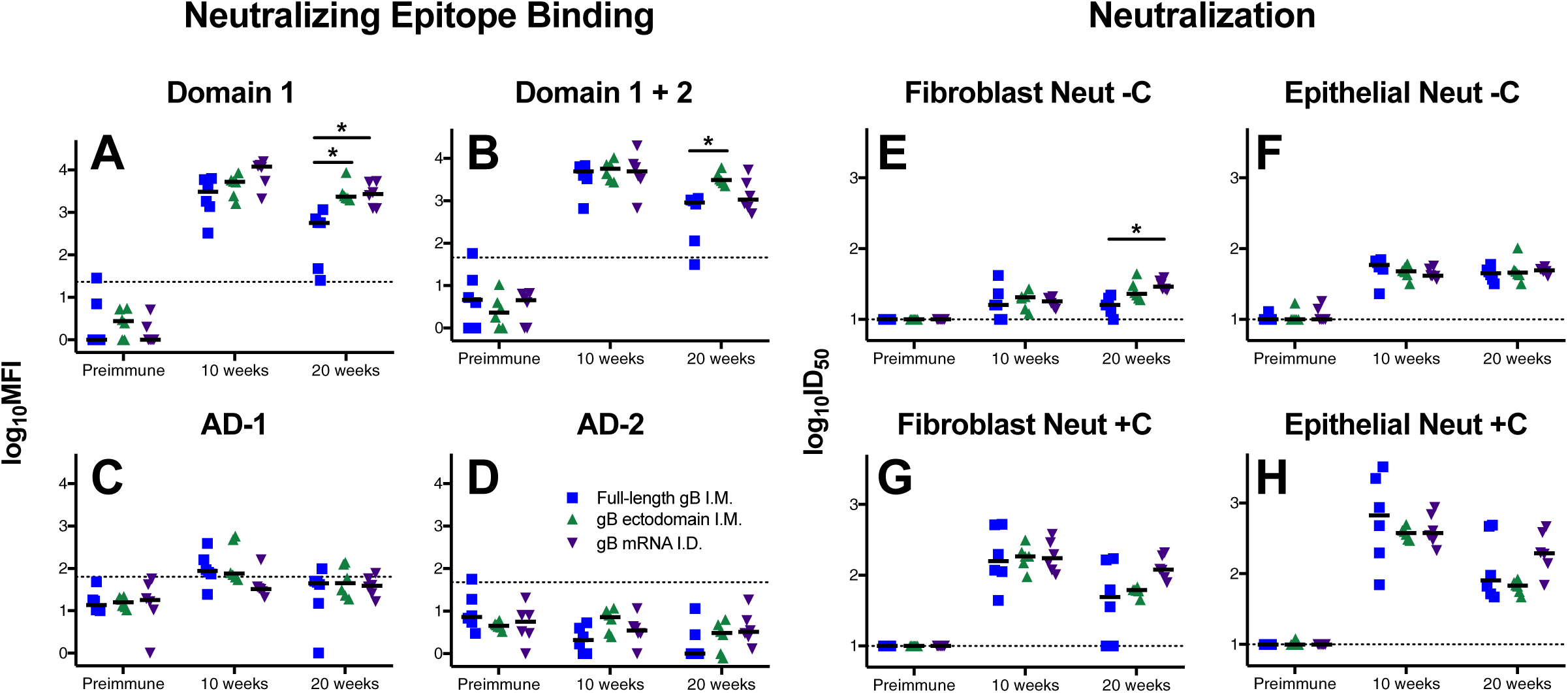
Enhanced durability of HCMV-neutralizing antibodies following gB mRNA-LNP vaccination. Vaccine-elicited binding to glycoprotein B neutralizing epitopes domain 1 (A), domain 1+2 (B), AD-1 (C), and AD-2 site 1 (D) was assessed by BAMA. Neutralization of heterologous virus AD169r in the absence (-C; E-F) and presence (+C; G-H) of purified rabbit complement on MRC-5 fibroblast cells (E,G) and ARPE epithelial cells (F,H). Full-length gB vaccinees are shown in blue, gB ectodomain in green, and gB mRNA-LNP in purple. Antibody responses were assessed for preimmune, 10 week (post boost 3, peak immunogenicity), and 20 week (necropsy) timepoints. Data points represent individual animals, with the line designating the median. Dotted black line indicates the mean preimmune response + 2 standard deviations. *p<0.05, Kruskal-Wallis + *post hoc* Mann-Whitney U test.

Furthermore, we assessed the magnitude of vaccine-elicited HCMV neutralization against heterologous (cross strain) AD169-revertant virus in fibroblast and epithelial cell lines, both in the presence and absence of purified rabbit complement (**Figure 4E-H**). Neutralization was enhanced in the presence of complement (**Figure 4G,H**) as previously noted [18]. All three vaccines had similar peak neutralization at 10 weeks (median AD169r ID_50_ in fibroblast at 10 weeks +C: FL = 158, ecto = 184, mRNA = 174). gB mRNA vaccinated rabbits had two-fold higher virus neutralization at 20 weeks, indicating superior response durability, though this distinction was not statistically significant (median AD169r ID_50_ in fibroblast at 20 weeks +C: FL = 49, ecto = 62, mRNA = 120, p=0.12, Kruskal-Wallis + *post hoc* Mann-Whitney U test). Notably, neutralizing responses measured in both fibroblast and epithelial cells were similar in magnitude, as might be expected for gB-specific antibodies. In addition to heterologous AD169r, we also measured neutralization of autologous (vaccine strain) Towne virus in fibroblast cells (**Figure S2A,B**), but identified little difference between vaccination groups.

### Vaccine-elicited engagement of F_c_γ receptors and non-neutralizing effector functions

We next investigated the ability of vaccine-elicited antibodies to engage F_c_γ receptors (specifically F_c_γRI, F_c_γRIIa, F_c_γRIIb, and F_c_γRIIIa) which is a prerequisite for F_c_-mediated effector functions (**Figure 5A-D**). Intriguingly, a single dose of gB mRNA vaccine (but not gB FL or gB ecto) resulted in the rapid development of antigen-specific IgG that could engage with F_c_ receptors (**Figure S3**), which might be due to robust induction of T follicular helper cells that facilitate B cell maturation and class switching [21]. This distinction was most notable for F_c_γRIIa and F_c_γRIIb (2 weeks median F_c_γRIIa MFI: FL = 0 vs. mRNA = 128, p=0.02, Kruskal-Wallis + *post hoc* Mann-Whitney U test). At peak immunogenicity (10 weeks), gB-specific antibody engagement of F_c_γ receptors was similar between vaccine groups. However, at the time of necropsy we generally noted enhanced median binding to F_c_γR’s among gB mRNA-vaccinated rabbits vs. gB FL, though this comparison was only significant for F_c_γRIIIa (20 weeks median F_c_γRIIIa MFI: FL = 3,929 vs. mRNA = 10,933, p=0.02, Kruskal-Wallis + *post hoc* Mann-Whitney U test).

**Figure 5.**
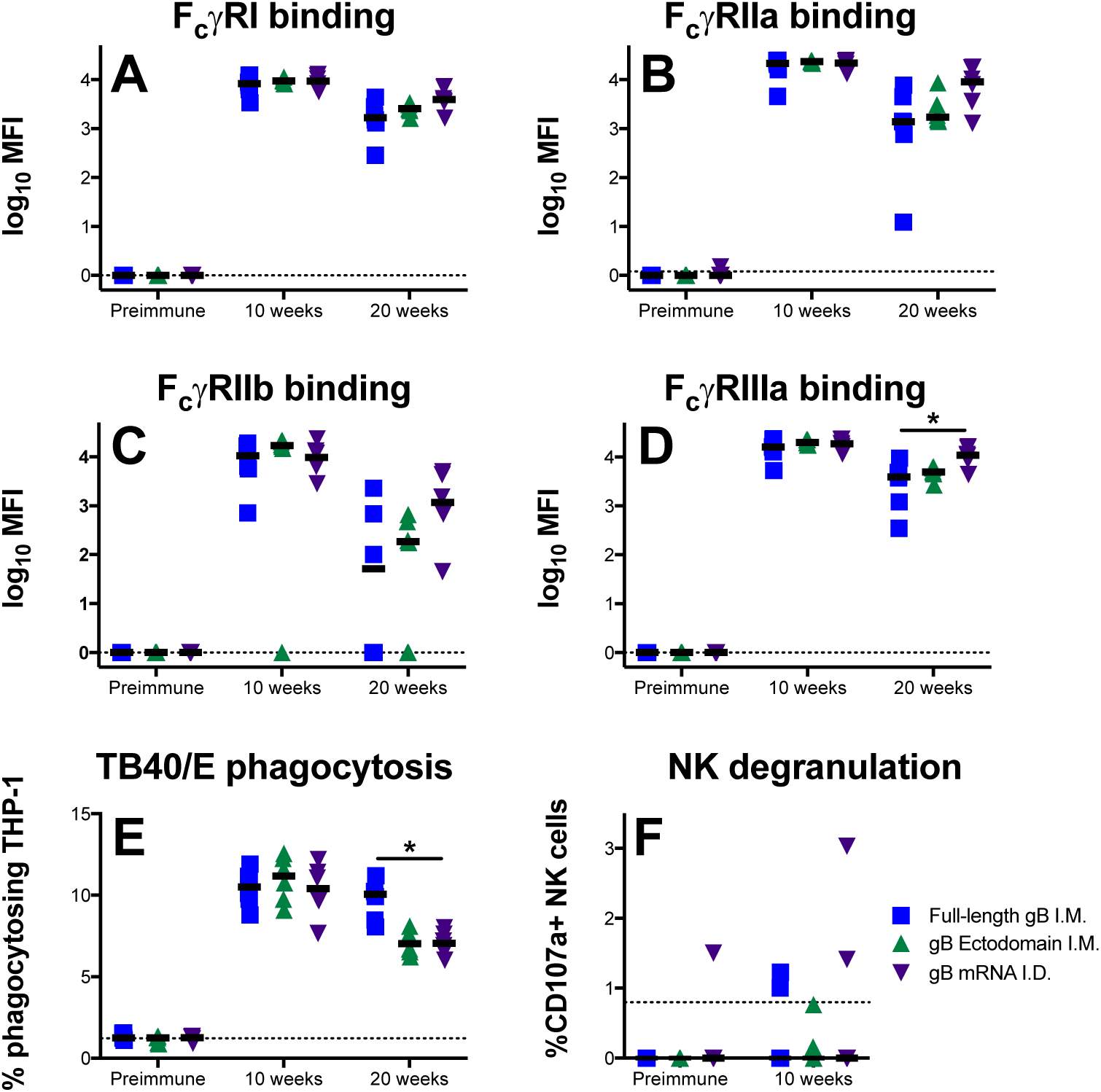
gB mRNA-LNP vaccination elicits long-lived gB-specific IgG-F_c_γ receptor engagement, though reduced durability of virion phagocytosis response. Vaccine-elicited IgG engagement of Fc**γ**RI (A), Fc**γ**RIIa (B), Fc**γ**RIIb (C), and Fc**γ**RIIIa (D) was assessed by BAMA. Phagocytosis of fluorophore-coupled whole HCMV (TB40/E) virions was measured by flow cytometry (E). Upregulation of CD107a on the surface of NK cells in the presence of HCMV-infected cells was assessed by flow cytometry as an approximation of ADCC-activity (F). Full-length gB vaccinees are shown in blue, gB ectodomain in green, and gB mRNA-LNP in purple. Antibody responses were assessed for preimmune, 10 week (post boost 3, peak immunogenicity), and 20 week (necropsy) timepoints. Data points represent individual animals, with the line designating the median. Dotted black line indicates the mean preimmune response + 2 standard deviations. *p<0.05, Kruskal-Wallis + *post hoc* Mann-Whitney U test.

We next investigated the magnitude of non-neutralizing, F_c_ effector functions mediated by vaccine-elicited antibodies. First, we measured antibody-dependent cellular phagocytosis (ADCP) of whole HCMV virions (TB40/E strain) (**Figure 5E**). We identified similar peak phagocytosis activity (10 weeks), but intriguingly we noted enhanced phagocytosis in gB FL vaccinees at 20 weeks indicating more robust phagocytosis durability in this group (median %phagocytosing cells: FL = 10.1% vs. mRNA = 7.1%, p<0.01, Kruskal-Wallis + *post hoc* Mann-Whitney U test). Next, we assessed NK cell degranulation activity (CD107a upregulation) when vaccine-elicited antibodies are incubated with HCMV-infected cells (**Figure 5F**), which we previously identified to approximate antibody-dependent cellular cytotoxicity (ADCC) for rabbits [17]. However, we were unable to measure NK degranulation activity for the majority of animals, suggesting that antibodies mediating this non-neutralizing effector function are not a dominant response elicited by gB FL, gB ecto, or gB mRNA vaccines.

### gB non-ectodomain-directed antibodies can mediate whole HCMV virion phagocytosis

We depleted gB ectodomain-specific IgG from vaccinated rabbit sera, and depletion was confirmed by ELISA against the same protein (**Figure 6A**). Furthermore, we established that AD-3-specific antibodies remained in gB ectodomain-depleted sera of gB FL vaccinees by measuring responses to an immunodominant AD-3 peptide (**Figure 6B).** Interestingly, ectodomain-depleted rabbit sera was able to mediate low-level HCMV virion phagocytosis (**Figure 6C**) (gB FL median % phagocytosing cells: preimmune = 0.96%, mock depleted = 2.91%, gB ecto depleted = 1.93%), suggesting that non-ectodomain epitopes (e.g. AD-3 or MPER) can be bound by circulating IgG that can subsequently mediate non-neutralizing effector functions. Furthermore, the magnitude of vaccine-elicited binding to AD-3 linear peptides of non-depleted plasma correlated strongly with phagocytosis mediated by gB ectodomain-depleted serum antibodies (**Figure 6D**; r = 0.75, p<0.001, Spearman-rank correlation).

**Figure 6.**
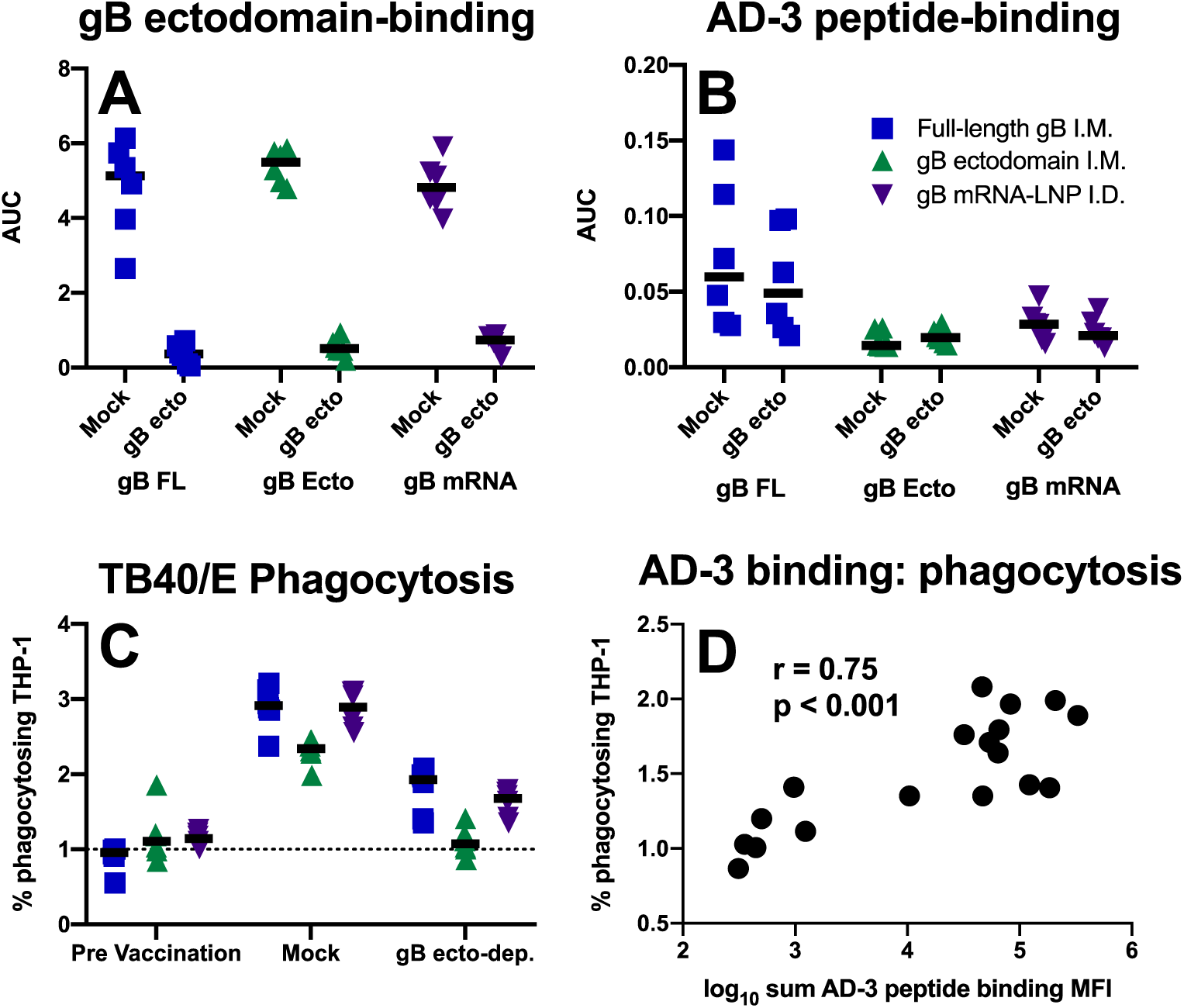
Phagocytosis-mediating antibodies directed against gB AD-3. Binding antibodies targeting gB ectodomain are not detectable in gB ectodomain-depleted sera (A), though antibodies targeting an immunodominant linear peptide within the AD-3 region (QDKGQKPNLLDRLRH) persist (B). ‘Mock’ = mock depleted sera using HIV-1 gp120, ‘gB ecto’ = gB ectodomain depleted sera. (C) Phagocytosis of whole HCMV (TB40/E) virions remained measurable by flow cytometry for samples depleted of ectodomain-targeting antibodies (i.e. with AD-3-specific antibodies remaining). (D) Spearman correlation of AD-3 peptide binding IgG magnitude with phagocytosis activity mediated by gB ectodomain-depleted serum antibodies. Full-length gB vaccinees are shown in blue, gB ectodomain in green, and gB mRNA-LNP in purple. Data points represent individual animals, with the line designating the median. Dotted black line indicates the mean preimmune response + 2 standard deviations.

### gB-specific T cell responses

Lastly, we investigated the magnitude of antigen-specific T cells by intracellular cytokine staining. Purified spleen mononuclear cells were either not stimulated, incubated with mitogen concanavalin A (ConA), or incubated with pooled gB peptides. Subsequently, the concentration of IFNγ^+^ live T cells identified for each group/treatment (**Figure 7A**). For animals from each vaccine group, ConA nonspecifically stimulated a population of T cells to produce IFNγ^+^ (all p<0.05, Friedman test + *post hoc* Wilcoxon matched pairs signed-rank test). When the percentage of unstimulated IFNγ^+^ T cells is subtracted from that of gB-stimulated IFNγ^+^ T cells (**Figure 7B**), we observed a modest gB-specific T cell response that is most pronounced in gB FL and gB mRNA vaccinated rabbits (median % gB-specific IFNγ^+^ T cells: FL = 0.22%, ecto = 0%, mRNA = 0.38%, p=ns, Kruskal-Wallis). In addition to spleen cells, we attempted to identify antigen-specific T cells in peripheral blood as well as mesenteric lymph nodes, but no IFNγ^+^ cells were observed using this method upon either ConA or gB peptide stimulation.

**Figure 7.**
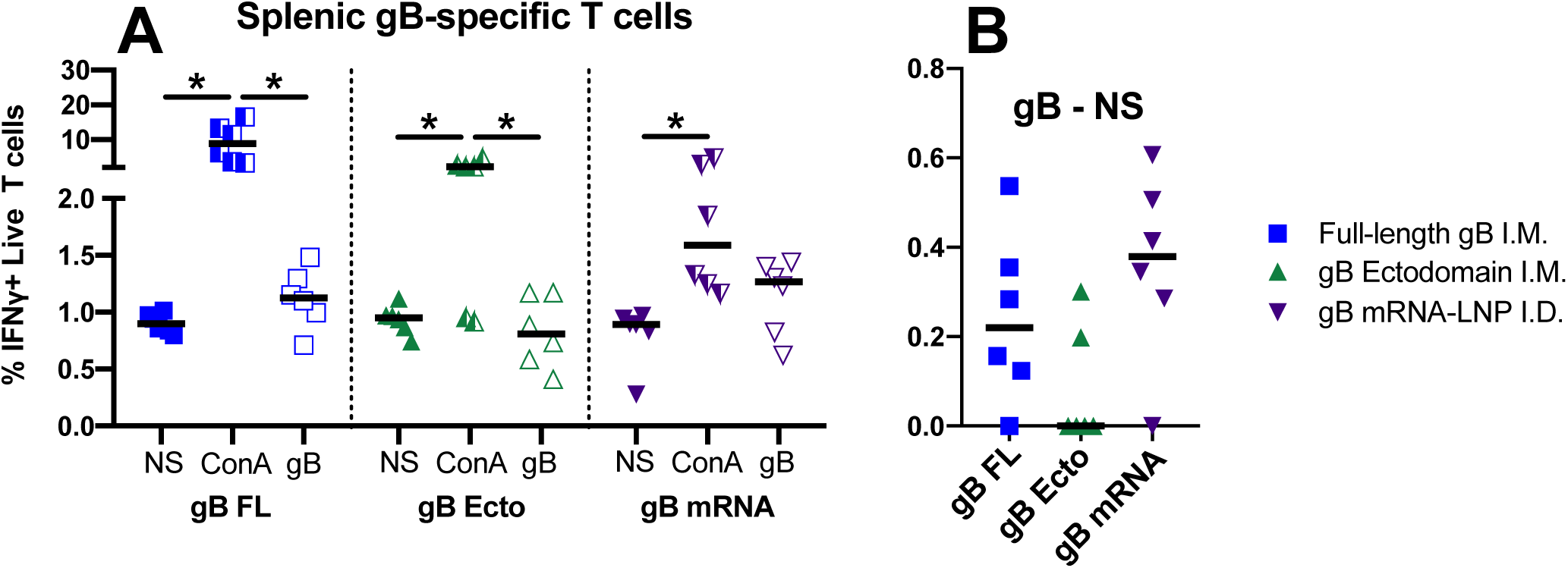
Full-length gB and gB mRNA-LNP vaccines elicit antigen-specific T cells in spleen of majority of vaccinees. (A) Splenic cells were either not stimulated (NS), incubated with Concanavalin A (ConA), or with a pool of gB peptides (gB), then stained for Ken-5 (rabbit pan T cell marker) and IFNγ. The percentage of live T cells that stained positive for IFNγ are plotted. (B) For each animal, the difference between the percentage of gB-stimulated and unstimulated cells is plotted. Full-length gB vaccinees are shown in blue, gB ectodomain in green, and gB mRNA-LNP in purple. Data points represent individual animals, with the line designating the median. *p<0.05, Friedman test + *post hoc* Wilcoxon matched pairs signed-rank test.

## Discussion

Glycoprotein B, a homotrimeric viral fusogen that is essential for entry into all cell types, has long been a leading HCMV vaccine candidate [9]. Yet over the past decade, following discovery that the most potent HCMV-neutralizing antibodies in human sera target the gH/gL/UL128-131A pentameric complex [22, 23], the focus has expanded from gB vaccine development. Nevertheless, it is important to recognize that the gB/MF59 protein subunit vaccine achieved partial moderate vaccine efficacy in preventing primary HCMV infection and seroconversion [12, 13] – a feat unparalleled in the HCMV vaccine field [24]. Furthermore, the gB/MF59 vaccine reduced viremia and demonstrated a protective benefit in transplant recipients [14]. Importantly, these partial successes were achieved without the elicitation of robust neutralizing antibody responses [16, 25]. In this investigation we sought to improve upon the gB immunogen antigenicity and vaccine/delivery platform, comparing novel vaccine immunogenicity head-to-head against the gB/MF59 vaccine in a preclinical model.

HCMV preclinical vaccine development is hindered by the fact that small animal models poorly represent host-pathogen biology and mechanisms of disease pathogenesis [26, 27]. We chose to test immunogenicity in rabbits because we previously demonstrated that vaccination can elicit antibodies with both neutralizing and non-neutralizing *in vitro* functionality [17]. In this study we were able to epitope-map and define the function of antibodies elicited by these three experimental vaccines, then compare to previously-reported immune correlates of protection (Nelson, *Journal of Infectious Diseases*, 2019, in press). HCMV-neutralizing IgG has previously been associated with reduced viral systemic dissemination [20, 28, 29]. Specifically, antibodies targeting gB AD-2 are correlated with reduced incidence of viremia and congenital disease [19, 20], though AD-2-specific antibodies were not elicited to any appreciable extent by the vaccines tested in this investigation (**Figure 4H**). Furthermore, non-neutralizing antibodies targeting gB and other surface glycoproteins are well described to have a protective role in preventing HCMV acquisition [16, 25], reducing viremia [16, 30], and blocking tissue-invasive replication [30, 31], though the precise mechanism remains unknown. Lastly, HCMV-specific CD4^+^ and CD8^+^ T cells have been widely implicated in reducing HCMV acquisition [32, 33], viral replication [26, 34-42], and the incidence of congenital/transplant-associated disease [43, 44].

Given the uncertainty regarding precise epitope specificities or immune effector functions protective against HCMV-associated disease, we regarded enhanced *durability* and *breadth* of immune responses as desirable attributes for vaccine development and the primary outcomes of interest in this study. Overall, we identified that the experimental vaccines (gB FL, gB ecto, and gB mRNA) elicited comparable magnitude binding and functional antibody responses at peak immunogenicity. Most distinction between vaccine-elicited immunogenicity was observed only at the latest timepoint (20 weeks – 12 weeks after final vaccine dose), signifying variable durability of elicited antibody responses. Nucleoside-modified mRNA-LNP vaccines have been well-described to produce sustained antigen presentation with robust T follicular helper cell and germinal center B cell stimulation, resulting in extraordinarily durable antibody responses after even a single vaccine dose in small and large animals [21, 45-47]. Indeed, we identified that the durability of gB mRNA vaccine-elicited responses exceeded that of gB FL for nearly all measured antibody binding/functional responses (**Table 1**; comparison statistically significant for: soluble and cell-associated gB binding, binding to gB domain 1 and domain 1+2, heterologous virus neutralization, and engagement of F_c_γRIIIa). Furthermore, we noted enhanced breadth of the peptide-binding immune response in gB mRNA-immunized rabbits, resulting in the targeting of unconventional/sub-dominant epitopes that are not observed in protein subunit vaccinees (**Figure 3, Figure S1**). mRNA-vaccinated rabbits appear to have a gB peptide-binding fingerprint that closely resembles that in seropositive individuals elicited by natural host infection [25]. Importantly, while HCMV gB mRNA vaccines have been tested previously in a preclinical model [48], this is the first such investigation to test a gB mRNA-based vaccine alongside the partially-effective gB/MF59 protein subunit vaccination and to directly compare the epitope specificity, durability, and neutralizing/non-neutralizing function of antibodies elicited by these two vaccine platforms.

**Table 1.** Summary of vaccine-elicited immune responses at week 20.

| Category | Median response magnitude | gB FL | gB ecto | gB mRNA |
| --- | --- | --- | --- | --- |
| **gB Binding** | gB FL binding (log<sub>10</sub> MFI) | 2.85 | 3.37 | **3.53\*** |
|  | gB ecto binding MFI (log<sub>10</sub> MFI) | 2.77 | 3.19 | **3.34\*** |
|  | Cell-assoc. gB binding (% cells) | 18.81 | 18.58 | **29.05\*** |
|  | RAI (gB FL target) | **0.93** | 0.90 | 0.78 |
| **Peptide binding<sup>+</sup>** | Total binding sum (log<sub>10</sub> MFI) | **5.25** | 4.74 | 5.04 |
|  | AD-3 binding sum (log<sub>10</sub> MFI) | **5.12** | 2.27 | 4.77 |
|  | # peptides bound (>100 MFI) | 44.5 | 28.5 | **85\*** |
| **Neut. Epitope Binding and Neutralization** | Domain I binding (log<sub>10</sub> MFI) | 2.76 | 3.37 | **3.44\*** |
|  | Domain I+II binding (log<sub>10</sub> MFI) | 2.96 | **3.49** | 3.03 |
|  | AD-1 binding (log<sub>10</sub> MFI) | NMR<sup>†</sup> | NMR | NMR |
|  | AD-2 binding (log<sub>10</sub> MFI) | NMR | NMR | NMR |
|  | AD169r Fibro Neut +C (log<sub>10</sub> ID<sub>50</sub>) | 1.69 | 1.79 | **2.08** |
|  | AD169 Epi Neut +C (log<sub>10</sub> ID<sub>50</sub>) | 1.91 | 1.83 | **2.29** |
| **Fcγ receptor binding and function** | FcγRI binding (log<sub>10</sub> MFI) | 3.22 | 3.41 | **3.59** |
|  | FcγRIIa binding (log<sub>10</sub> MFI) | 3.14 | 3.24 | **3.29** |
|  | FcγRIIb binding (log<sub>10</sub> MFI) | 1.71 | 2.27 | **3.07** |
|  | FcγRIIIa binding (log<sub>10</sub> MFI) | 3.59 | 3.69 | **4.04\*** |
|  | TB40/E virion phagocytosis (% cells) | **10.06\*** | 7.04 | 7.06 |
|  | NK degranulation (% cells) | NMR | NMR | NMR |
| **T cells** | Splenic gB-specific T cells (BS, % live T) | 0.22 | NMR | **0.38** |

\* p<0.05, Kruskal-Wallis + *post hoc* Mann-Whitney U test (compared to gB FL)
<sup>+</sup> Peptide binding using week 10 sera
<sup>†</sup> NMR = no measurable response above baseline levels

In this investigation, we specifically focused on the implications of the gB/MF59-elicited immune-dominant response directed against gB AD-3 [25]. We hypothesized the absence of neutralizing antibodies in gB/MF59 vaccinees may be attributable to the dominant responses against the AD-3 ‘decoy epitope’, which diverted antibody targeting away from ‘more functional’ epitopes [24]. We directly tested this hypothesis with our gB ecto group by excluding AD-3 from the immunogen. Intriguingly, we failed to see any consistent increase in the magnitude of functional neutralizing/non-neutralizing antibodies in gB ecto vaccinated rabbits, suggesting that inclusion of the AD-3 epitope in the vaccine immunogen does not hinder the development of more functional antibodies. Is it therefore possible that AD-3 directed antibodies have any functional or protective role that might account for the 50% vaccine efficacy observed in gB/MF59 vaccinees? In this study we noted that AD-3-specific antibodies can mediate non-neutralizing antibody effector functions including whole virion phagocytosis (**Figure 6**). Consequently, the high-magnitude AD-3 response in gB FL-vaccinated rabbits likely accounts for our observation of robust and durable phagocytosis activity in these animals (**Figure 5E**).

A limitation to this study is dissimilarity in vaccine dose and route of delivery between comparison groups. Protein subunit vaccines (gB FL, gB ecto) were given I.M. at a dose of 20μg, mimicking the protocol for gB/MF59 immunized humans in the partially-efficacious clinical trials [12, 13, 24]. In contrast, 50μg of the gB nucleoside-modified mRNA-LNP was administered I.D, which was selected because this delivery method enhances protein expression *in vivo* [49]. Depending on the antigen and vaccine formulation, intradermal vaccination may be as much as 10 times more potent than intramuscular dosing [50]. Therefore, we cannot rule out the possibility that our results of enhanced breadth and durability in gB mRNA-immunized rabbits are dose/method dependent – that a higher dose or different delivery method of gB FL protein might not have achieved similar results to gB mRNA-LNP vaccinated rabbits. Furthermore, while rabbits provide an excellent model to study vaccine immunogenicity, this investigation was restricted by the rabbit immunologic toolbox. We were able to measure F_c_γ receptor engagement in this study, though lacked the ability to identify the mechanism behind variable F_c_γ receptor engagement (e.g. IgG subclass or F_c_ glycosylation). Furthermore, while we were able to identify gB-specific T cells in spleen, we were unable to: 1) identify antigen-specific T cells in peripheral blood, and 2) parse out T cell subsets (CD4^+^, CD8^+^, etc). Nevertheless, the results described are sufficient justification for subsequent testing of the gB mRNA-LNP vaccine in nonhuman primate preclinical challenge models and/or human clinical trials.

This comparison of immune responses elicited by next-generation HCMV vaccines, head-to-head against gB/MF59 immunization (gB FL), provides a basis for rational gB vaccine development efforts. First, we have demonstrated that gB nucleoside-modified mRNA-LNP immunization improves the durability of gB-binding and functional antibody responses and enhances the breadth of the gB-specific antibody repertoire. Additionally, we note that gB ectodomain immunization did not elicit antibody responses that were functionally superior to gB FL, suggesting that the immunodominant AD-3 response does not interfere with the development of functional antibodies. While we await well-validated immune correlates of protection to inform HCMV vaccine development efforts, the gB mRNA-LNP vaccine clearly had enhanced long-term immunogenicity and response breadth compared with gB FL and gB ecto vaccines. Therefore, we propose subsequent testing of this vaccine platform in nonhuman primate challenge models as well as human immunogenicity trials to rigorously interrogate its ability to elicit immune factors protective against congenital HCMV infection and transplant-associated disease.

## Methods

### gB ectodomain immunogen production

The sequence encoding the ectodomain segment (amino acid residues 1-696) of Towne strain (GenBank accessioning# FJ616285.1) HCMV glycoprotein B (gB) was tagged at the 3’ end with a polyhistidine tag, and the furin cleavage site at residue 457 mutated from ‘RTKR’ to ‘STKS’. The nucleotide sequence was codon-optimized for mammalian cells, then cloned into pcDNA3.1(+) mammalian expression vector (Invitrogen) via *BamHI* site at the 5’ end and *EcoRI* site at the 3’ end. Subsequently, the plasmid was transiently transfected into Expi293i cells using ExpiFectamine 293 transfection reagents (ThermoFisher Scientific) according to the manufacturer’s instructions. Culture supernatant was harvested after 5 days of incubation at 37 °C and 8% CO_2_, then purified using Nickel-NTA resin (ThermoFisher Scientific). Purity, identity, and correct molecular weight were confirmed by Western blot using monoclonal antibodies specific for gB AD-2, gB domain 1, and gB domain 2 (described in [15]), followed AP-conjugated anti-human IgG (Sigma-Aldrich). Finally, the protein was tested for the presence of endotoxin using the Pierce LAL chromogenic endotoxin quantitation kit (ThermoFisher Scientific). gB protein ectodomain aliquots were stored at −80°C at a concentration of ∼1 μg/μl, then thawed <60 minutes prior to injection.

### gB mRNA production and formulation into lipid nanoparticles

The modified mRNA encoding HCMV gB (Towne strain, GenBank accessioning# FJ616285.1) was produced as previously described [51] using T7 RNA polymerase (Megascript, Ambion) on codon-optimized [52] linearized plasmid (sequence is available upon request). The mRNA was transcribed to contain 101 nucleotide-long poly(A) tail. To generate modified nucleoside-containing mRNA, m1ψ-5’-triphosphate (TriLink) was used instead of UTP. The mRNA was then capped using the m7G capping kit with 2’-*O*-methyltransferase (ScriptCap, CellScript). The mRNA was purified by Fast Protein Liquid Chromatography (FPLC) (Akta Purifier, GE Healthcare), as described [53] and analyzed by electrophoresis using denaturing or native agarose gels, and stored at −20 °C. The FPLC-purified m1ψ-containing HCMV gB mRNA and poly(C) RNA (Sigma) were encapsulated in LNPs using a self-assembly process in which an aqueous solution of mRNA at pH=4.0 is rapidly mixed with a solution of lipids dissolved in ethanol [54]. LNPs used in this study were similar in composition to those described previously [54, 55], which contain an ionizable cationic lipid (proprietary to Acuitas)/phosphatidylcholine/cholesterol/PEG-lipid (50:10:38.5:1.5 mol/mol) and were encapsulated at an RNA to total lipid ratio of ∼0.05 (wt/wt). They had a diameter of ∼80 nm as measured by dynamic light scattering using a Zetasizer Nano ZS (Malvern Instruments Ltd) instrument. mRNA-LNP formulations were stored at −80°C at a concentration of mRNA of ∼1 μg/μl, then thawed <60 minutes prior to injection.

### Animal care and sample collection

Juvenile New Zealand White rabbits (approximately 10 weeks of age), were purchased from Robinson Services Inc (Mocksville, NC) and housed at Duke University. For blood collections, animals were sedated with 1mg/kg subcutaneous acepromazine and topical 1% lidocaine applied to the ears. EDTA-anticoagulated blood was collected via auricular venipuncture. Plasma was separated from whole blood by centrifugation, and PBMCs were isolated by density gradient centrifugation using Lympholyte cell separation media (Cedarlane laboratories). Animals were euthanized using 0.5mL of subcutaneously-injected xylazine (100mg/mL) + ketamine (500mg/mL) mixed together in a 1:5 ratio, followed by 0.5mL of intracardiac Euthasol (pentobarbital sodium + phenytoin sodium). Lymphocytes were isolated from spleen and mesenteric lymph nodes by manual tissue disruption and crushing through a 100μm cell strainer, followed by density gradient centrifugation with Lympholyte cell separation media.

### Animal vaccination

Three groups of juvenile New Zealand White rabbits (n=6) were given different vaccines: 1) 50μg intramuscular full-length gB protein (generous gift of Sanofi Pasteur) combined 1:1 v/v with MF59-like squalene adjuvant AddaVax (Invivogen), 2) 50μg intramuscular gB ectodomain protein combined 1:1 v/v with AddaVax, or 3) 50μg intradermal nucleoside-modified gB mRNA packaged in lipid nanoparticles. Vaccine doses were administered monthly for three consecutive months. For intramuscular (I.M.) protein subunit vaccine administration, rabbits were sedated with 1mg/kg subcutaneous acepromazine then injected with the vaccine dose in the rear thigh (alternating sides between monthly doses). For intradermal (I.D.) mRNA-LNP vaccine administration, rabbits were sedated with 1mg/kg subcutaneous acepromazine as well as 2% inhaled isofluorane, and the saddle of the rabbit shaved. Chilled, sterile PBS was added to each vaccine dose to a total volume of 300 μL, the diluted vaccine divided into 6 equal fractions of 50μL each, then the rabbit saddle was injected intradermally with 3 injections on each side of the spine (injection sites varied between monthly doses).

### Cell culture

Human retinal pigment epithelial (ARPE-19) cells (ATCC) were maintained for a maximum of 35 passages in Dulbecco’s modified Eagle medium-12 (DMEM-F12) supplemented with 10% FCS, 2mM L-glutamine, 1mM sodium pyruvate, 50 U/mL penicillin, 50 μg/mL streptomycin and gentamicin, and 1% epithelial growth cell supplement (ScienCell). Human lung (MRC-5) fibroblasts (ATCC) were maintained for a maximum of 20 passages in DMEM containing 20% FCS, 50 U/mL penicillin, and 50 μg/mL streptomycin. Human epithelial kidney (HEK293T) cells (ATCC) were maintained for a maximum of 35 passages in DMEM containing 10% FCS, 25mM HEPES buffer, 50 U/mL penicillin, and 50 μg/mL streptomycin. Human monocyte (THP-1) cells (ATCC) were maintained for a maximum of 35 passages in RPMI-1640 medium containing 10% FCS. All cell lines were tested for the presence of mycoplasma biannually.

### Virus growth

AD169 revertant virus (AD169r; a gift from Merck) [56], and BADUL131a virus [57] stocks were propagated on ARPE cells in T75 culture flasks. Towne virus (ATCC) was propagated on MRC-5 cells in T75 culture flasks. Supernatant containing cell-free virus was collected when 90% of cells showed cytopathic effects, then cleared of cell debris by low-speed centrifugation before passage through a 0.45-μm filter. Viral infections of ARPE-19 cells were carried out in similar media but contained only 5% FCS and lacking cell growth supplement.

### Binding antibody multiplex assay (BAMA)

Antibody responses against gB full-length protein, gB ectodomain, and gB epitopes (creation described in [14, 20]) were assessed by multiplex ELISA. In brief, carboxylated fluorescent beads (Luminex) were covalently coupled to purified HCMV antigens and subsequently incubated with maternal plasma in assay diluent (phosphate-buffered saline, 5% normal goat serum, 0.05% Tween 20, and 1% Blotto milk, 0.5% polyvinyl alcohol, and 0.8% polyvinylpyrrolidone). The antigen panel included full-length gB (courtesy of Sanofi-Pasteur), gB ectodomain protein, gB domain 1, gB domain 2, gB domain 1+2, gB AD-1 (myBiosource), and biotinylated linear gB AD-2 (biotin-NETIYNTTLKYGD). HCMV glycoprotein– specific antibodies were detected with phycoerythrin-conjugated goat anti-human IgG (2 μg/mL, Southern Biotech). Beads were washed and acquired on a Bio-Plex 200 instrument (Bio-Rad), and results were expressed as mean fluorescence intensity. A panel of pre-vaccination time points was tested to determine nonspecific baseline levels of binding. Minimal background activity was observed, so the threshold for positivity for each antigen was set at the mean value of negative control sera to each antigen + 3 standard deviations. Blank beads were used in all assays to account for nonspecific binding. All assays included tracking of HCMV immunoglobulin (Cytogam – CSL Behring) standard by Levy-Jennings charts. The preset assay criteria for sample reporting were coefficient of variation per duplicate values of ≤20% for each sample and ≥100 beads counted per sample. All samples were analyzed at the same dilution for each antigen: full-length gB, gB ectodomain, gB domain 1, gB domain 2, and gB domain 1+2 were assessed at a 1:500 dilution; gB AD-1 and gB AD-2 were assessed at a 1:50 dilution. These dilutions were predetermined to be within the linear range of the assay based on testing serial dilutions of a small subset of plasma samples.

### F_c_γ receptor engagement

The binding of vaccine-elicited serum antibodies to F_c_γ receptors was characterized using a multiplex F_c_γ receptor BAMA assay, employing the reagents and QC methods described above. In brief, HCMV gB (full-length) was covalently coupled to streptavidin (Rockland)-coupled fluorescent beads (Luminex). Sera samples were diluted 1:500, then incubated in duplicate in a 96-well microplate with gB-streptavidin-coupled beads, then washed and incubated with one of the following biotinylated F_c_γ receptor tetramers: F_c_γRIa, F_c_γRIIa (clone H131), F_c_γRIIb, and F_c_γRIIIa (clone V158) (F_c_γ receptor proteins courtesy of Dr. Kevin Saunders). F_c_γ receptor engagement was detected using mouse anti-human IgG-PE (myBioSource) followed by a final wash. Data were acquired on a BioPlex-200 (Luminex).

### gB-transfected cell binding

Binding of vaccine-elicited antibodies to trimeric gB expressed on cell membranes was assessed by flow cytometry as previously [15]. Briefly, HEK293T cells were grown overnight to ∼50% confluency, then co-transfected using Effectine transfection reagent (Qiagen) with a GFP-expressing plasmid (gift of Maria Blasi, Duke University) and a second plasmid encoding the full-length Towne strain gB (SinoBiological). Transfected cells were incubated for 2 days at 37°C and 5% CO_2_, washed with DPBS (Gibco), then removed from the flask using enzyme free cell-dissociation buffer (ThermoFisher Scientific). Cells were washed in wash buffer (DPBS + 1% FBS), then 100,000 live cells were added to each well of a 96-well V-bottom plate (Corning). After centrifugation (1200 x g, 5 minutes), cells were re-suspended in 1:6250 diluted sera samples and incubated for 2 hours at 37°C and 5% CO_2_. Next, cells were washed and stained with LIVE/DEAD Aqua Dead Cell Stain Kit (ThermoFisher Scientific) diluted 1:1000 for 20 minutes at RT. Afterwards, cells were washed, then re-suspended in PE-conjugated goat anti-human IgG Fc (eBioscience) diluted 1:200 in wash buffer then incubated for 25 minutes at 4°C. Following two additional wash steps, cells were re-suspended and fixed in DPBS + 1% formalin. Events were acquired on LSR Fortessa machine (BD biosciences) using the high-throughput sampler (HTS). The % PE+ cells was calculated from the live, GFP+ cell population and reported for each sample. Background binding of each plasma sample was corrected for using cells transfected with the GFP-expressing plasmid alone.

### Avidity ELISA

384-well ELISA plates (Corning) were coated overnight at 4°C with 30 ng full-length gB per well, then blocked with assay diluent (1x PBS containing 4% whey, 15% normal goat serum, and 0.5% Tween-20). Three-fold dilutions of sera were then added to the plate,, then duplicate wells were treated for 5 min with either 7M urea or 1x PBS following sera incubation. Finally, bound IgG was detected with a horseradish peroxidase (HRP)-conjugated polyclonal goat anti-monkey IgG (Rockland), and developed using the SureBlue Reserve tetramethylbenzidine (TMB) peroxidase substrate (KPL). Sera dilutions that resulted in an OD value between 0.6 and 1.2 in the absence of urea treatment (dilution range = 1:30-1:1000) were used to determine the relative avidity index (RAI). Indexes were calculated as the OD ratio of urea:PBS treated wells.

### Glycoprotein B peptide microarray

Binding to gB linear peptides was assessed as previously [15]. In brief, 15-mer peptides covering the entire gB open reading frame (Towne strain), and overlapping with neighboring peptides by 10 residues (total of 188 peptides) were synthesized and printed to a PepStar multiwell array (JPT Peptide) in triplicate. Microarray binding was performed manually using individual slides immobilized in the ArraySlide 24-4 chamber (JPT Peptide). First, arrays were blocked with blocking buffer (PBS containing 1% milk blotto, 5% NGS, and 0.05% Tween20), incubated first with sera diluted 1:250 in blocking buffer, and secondly with anti-human IgG conjugated to AF647 (Jackson ImmunoResearch) diluted in blocking buffer (0.75 μg/mL). Arrays were washed in wash buffer (1x TBS buffer + 0.1% Tween) between steps using an automated plate washer (BioTec ELX50). To measure fluorescence, arrays were scanned at a wavelength of 635 nm using an InnoScan 710 device (Innopsys) at a PMT setting of 580 and 100% laser power. Images were analyzed using Mapix software (Innopsys), and reviewed manually for accurate automated peptide identification. Binding intensity of sera to each peptide was corrected with the surrounding background fluorescence. Median fluorescent intensity of each of the 3 replicates was reported.

### Neutralization

The neutralization titers of patient sera were measured by both a high-throughput immunofluorescence assay as previously described [15]. Briefly, MRC-5 cells were seeded into 96-well flat-bottom plates and incubated for 2 days at 37°C and 5% CO2 to achieve 100% confluency. Once confluent, 3-fold dilutions (1:10–1:30,000) of heat-inactivated rabbit sera in infection media were incubated with an MOI=1.0 of Towne (ATCC) or AD169r (Merck Laboratories) virus stock in a total volume of 50 μL for 45 minutes at 37°C. For complement neutralization assays, plasma/virus was diluted in infection media containing rabbit complement (Cedarlane Laboratories) at a final dilution of 1:4. Immune complexes were added in duplicate to wells containing MRC-5 cells, then subsequently incubated for 18 hours at 37°C. Infected cells were then fixed for 10 minutes with 3.7% paraformaldehyde, permeabilized for 10 minutes with Triton × 100, and subsequently processed for immunofluorescence with mouse anti-HCMV IE-1 monoclonal antibody (MAB810, Millipore) followed by goat anti-mouse IgG-AlexaFluor488 (Millipore) and DAPI nuclear stain. Total cells and AF488+ infected cells per well were counted on a Cellomics Arrayscan (ThermoFisher Scientific). Neutralization titers (ID_50_) were calculated according to the method of Reed and Muench using the plasma dilution that resulted in a 50% reduction in the percentage of infected cells compared to control wells infected with virus only.

### Whole HCMV virion phagocytosis

The ability of vaccine-elicited antibodies to facilitate phagocytosis of whole HCMV virions was assessed as previously [15]. Briefly, 10^7^ PFU of concentrated, sucrose gradient-purified HCMV TB40/E-mCherry virus was buffer exchanged with 1x PBS and sodium bicarbonate (0.1M final concentration), then AF647 NHS ester (Invitrogen) added for direct viral conjugation at room temperature for 1 hour with constant agitation. The reaction was quenched with 1 M Tris-HCl, pH 8.0, then the labelled virus was diluted 25x in wash buffer (PBS + 0.1% FBS). Sera samples were diluted 1:10 in wash buffer, then 10 μL of diluted sera combined with 10 μL of diluted, fluorophore-conjugated virus in a round-bottom, 96-well plate (Corning) and allowed to incubate at 37°C for 2 hours. Following this incubation step, 25,000 THP-1 cells were added to each well, suspended in 200 μL primary growth media. Plates were centrifuged at 1200 xg and 4°C for 1 hour in a spinoculation step, then incubated at 37°C for an additional hour. Cells were re-suspended and transferred to a 96-well V-bottom plate, then washed twice prior to fixing in 100μL DPBS + 1% formalin. Events were acquired on LSR Fortessa machine (BD biosciences) using the HTS. The % AF647+ cells was calculated from the full THP-1 cell population and reported for each sample. A cutoff for sample positivity was defined as the mean value of pre-vaccination sera (n=18) + 2 standard deviations.

### Natural killer (NK) cell CD107a degranulation assay

Cell-surface expression of CD107a was used as a marker for NK cell degranulation, which we have previously shown to have good agreement with antibody-dependent cellular cytotoxicity activity for rabbit sera [17]. MRC-5 cells were infected with BadrUL131-Y4 with a MOI of 1.0 for 48 hours at 37°C, at 4×10^4^ cells/well in 96-well flat-bottom tissue culture plates. Following incubation, supernatant was removed and the infected cell monolayers were washed once with RMPI 1640 containing 10% FBS, HEPES, Pen-Strep-L-Glut, Gentamicin (R10 media) before addition of NK cells. Primary human NK cells were isolated from peripheral blood mononuclear cells (PBMC) after overnight rest in R10 media with 10ng/mL IL-15 (Miltenyi Biotech) by depletion of magnetically labeled cells (Human NK cell isolation kit, Miltenyi Biotech). 5×10^4^ live NK cells were added to each well containing HCMV-infected MRC-5 cell monolayers. Plasma samples were diluted in R10 and added to the cells at a final dilution of 1:50 in duplicate. Brefeldin A (GolgiPlug, 1 μl/ml, BD Biosciences), monensin (GolgiStop, 4μl/6mL, BD Biosciences), and CD107a-FITC (BD Biosciences, clone H4A3) were added to each well and the plates were incubated for 6 hours at 37oC in a humidified 5% CO_2_ incubator. NK cells were then gently resuspended, taking care not to disturb the MRC-5 cell monolayer, and the NK containing supernatant was collected and transferred to 96-well V-bottom plates. The recovered NK cells were washed with PBS, and stained with LIVE/DEAD Aqua Dead Cell Stain at a 1:1000 dilution for 20 minutes at room temperature. The cells were then washed with 1%FBS PBS and stained for 20 minutes at room temperature with the following panel of fluorescently conjugated antibodies diluted in 1%FBS PBS: CD56-PECy7 (BD Biosciences, clone NCAM16.2), CD16-PacBlue (BD Biosciences, clone 3G8), and CD69-BV785 (BioLegend, Clone FN50). The cells were then washed twice and re-suspended in 1% paraformaldehyde fixative for flow cytometric analysis. Data analysis was performed using FlowJo software (v9.9.6). Data is reported as the % of CD107a positive live NK cells (singlets, lymphocytes, aqua blue^-^, CD56^+^ and/or CD16^+^, CD107a^+^). CD69 was not used in the final analysis due to the low frequency of CD107a+ responses. Parallel assays were performed with uninfected MRC-5 cells as a control for identification of non-CMV specific responses, and final data are presented after subtraction of background activity observed against uninfected cells.

### Soluble protein bead coupling and antibody depletion

Cyanogen bromide-activated (CNBr-activated) sepharose beads (GE Healthcare) were rehydrated with 1 mM HCl then suspended in coupling buffer (0.1 M NaHCO_3_ + 0.5 M NaCl, pH 8.3). Every 100 μL of bead slurry was combined with 200 μg of soluble gB ectodomain (post-fusion conformation, courtesy of Jason McLellan, University of Texas, Austin). Coupling proceeded for 12 hours at 4°C on an inversion rotator. Excess soluble protein was washed off with 5 column volumes of coupling buffer. Unbound CNBr active groups were blocked with quenching buffer (0.1 M Tris HCl pH 8.0) for 2 hours at room temperature. Protein conjugated beads were washed with 3 cycles of alternating pH 0.1 M acetic acid, pH 4.0 and 0.1 M Tris HCl pH 8. For depletion of gB post fusion specific antibodies, 100 μg of protein-coupled bead slurry was loaded into a spin microelution column (Pierce, TFS). 400 μL of filtered (SpinX) 1:50 diluted plasma from vaccinated rabbits was then added to each column. Plasma was centrifuged through the column 10 times with bound IgG elution (0.2 M glycine elution buffer, pH 2.5) and bead recalibration (3 cycles of alternating pH washes) after spins 5 and 10. Adequate specific depletion was confirmed by ELISA against the depleted protein. Mock depleted samples underwent an identical depletion procedure in the presence of HIV-1 gp120 conjugated CNBr beads.

### Splenic T cell intracellular cytokine staining

Primary spleen cells were thawed in RPMI + 10% FBS with benzonase (50 U/mL) then measured for count and viability on Muse Cell Counter (Luminex). Cells were coincubated in duplicate with Con A (5ug/mL), HCMV UL155 (250ng/mL. JPT), or media for 20-24 hours at 37 degrees C. Samples were stained for rabbit pan-T cell marker KEN-5 (SCBT) and live/dead (Invitrogen), then fixed and permeabilized using BD Cytofix/Cytoperm according to the manufacturer’s instructions. Cells were stained for rabbit IFN-γ (mAb Tech). Flow cytometry was performed on a BD LSR II, and data was analyzed in Flow Jo v10. Gating strategy shown in **Figure S4**.

### Statistical analysis

Nonparametric tests were utilized because of the small group sizes (n=6 per group). Furthermore, the 20 week timepoint was employed for all statistical comparisons between vaccination groups. Magnitude of immune responses between the 3 vaccine groups were compared first by Kruskal-Wallis test. If p<0.05, *post hoc* Mann-Whitney U test was conducted for gB ecto and gB mRNA compared with gB FL. All statistical tests were carried out using the R statistical interface (version 3.3.1, www.r-project.org) and were two-tailed.

### Ethics statement

Animals were maintained in accordance with the American Association for Accreditation of Laboratory Animal Care standards and *The Guide for the Care and Use of Laboratory Animals* [58]. Efforts were made to minimize stress and provide enrichment opportunities when possible (social housing when possible, objects to manipulate in cage, varied food supplements, interaction with caregivers and research staff). All protocols were reviewed and approved by the Duke University Animal Care and Use Committee (IACUC) prior to the initiation of the study (protocol #A314-15-12).

## Author contributions

C.S.N. and S.R.P. designed research; C.S.N., J.A.J., N.P., H.R., M.G., and W.E. performed research; C.S.N. analyzed data; D.W., J.M., and J.P. contributed reagents and expertise; C.S.N. and S.R.P. wrote the paper.

## Acknowledgements

The authors would like to thank Diego Zapata and the Duke Division of Laboratory and Animal Resources for assistance with rabbit handling. Furthermore, we acknowledge Sanofi-Pasteur, Merck, and Trellis Biosciences for the generous gift of research materials. This work was supported by: NIH/NIAID R21 to S.R.P (R21AI136556) and NIH/NICHD F30 grant to C.S.N (F30HD089577). The funders had no role in study design, data collection and interpretation, decision to publish, or the preparation of this manuscript. The content is solely the responsibility of the authors and does not necessarily represent the official views of the National Institutes of Health.

## Supporting Information Legends

**Figure S1. Enhanced breadth of gB peptide-binding responses elicited by gB mRNA-LNP vaccination.** (A) Sum of linear gB peptide binding IgG magnitude at peak immunogenicity (week 10) to peptides within known antigenic epitopes – AD-1, AD-2 site 1, AD-2 site 2, AD-3, AD-4 (Domain 2), AD-5 (Domain 1), and the furin protease cleavage site. Binding to linear peptides outside these known antigenic regions is denoted ‘other’. Full-length gB vaccinees are shown in blue, gB ectodomain in green, and gB mRNA-LNP in purple. Data points represent individual animals, with the line designating the median. *p<0.05, Kruskal-Wallis + posthoc Mann-Whitney U test (B-D).

**Figure S2. No difference in durability of autologous neutralization between vaccination groups.** (A,B) Neutralization of Towne autologous virus in the absence (-C; A) and presence (+C; B) of purified rabbit complement on MRC-5 fibroblast cells. Full-length gB vaccinees are shown in blue, gB ectodomain in green, and gB mRNA-LNP in purple. Data points represent individual animals, with the line designating the median. Dotted black line indicates the mean preimmune response + 2 standard deviations.

**Figure S3. Rapid induction of gB-specific IgG that engage F**_**c**_***γ* receptor following a single mRNA vaccine dose.** Vaccine-elicited IgG engagement of Fc*γ*RI (A), Fc*γ*RIIa (B), Fc*γ*RIIb (C), and Fc*γ*RIIIa (D) was assessed by BAMA. Antibody responses were assessed for preimmune and 2 week (post boost 1) timepoints. Full-length gB vaccinees are shown in blue, gB ectodomain in green, and gB mRNA-LNP in purple. Data points represent individual animals, with the line designating the median. Dotted black line indicates the mean preimmune response + 2 standard deviations. *p<0.05, Kruskal-Wallis + posthoc Mann-Whitney U test.

**Figure S4. Example flow cytometry gating scheme for rabbit splenic T cells.** Spleen cells shown from ConA-stimulated, full-length gB I.M. vaccinated rabbit. (A) Cells were selected using forward and side scatter. (B) Live rabbit T cells were identified using AQUA live/dead stain (Invitrogen) as well as Ken-5 (pan-T cell marker) specific antibody (AF647). (C) IFN*γ*+ T cell subpopulation identified by intracellular cytokine staining (AF488).

